# Impact of type 2 diabetes on cardiorespiratory function and exercise performance

**DOI:** 10.1101/073205

**Authors:** Joanie Caron, Gregory R. duManoir, Lawrence Labrecque, Audrey Chouinard, Annie Ferland, Paul Poirier, Sylvie Legault, Patrice Brassard

**Affiliations:** Department of Kinesiology, Faculty of Medicine, Université Laval, Québec, Canada; Centre de recherche de l’Institut universitaire de cardiologie et de pneumologie de Québec, Québec, Canada; Faculty of Pharmacy, Université Laval, Québec, QC, Canada; School of Health & Exercise Sciences, University of British Columbia Okanagan, Kelowna, British Columbia, Canada

**Keywords:** maximal oxygen uptake, oxygen uptake kinetics, heart rate kinetics, type 2 diabetes

## Abstract

The aim of this study was to examine the impact of well-controlled uncomplicated type 2 diabetes (T2D) on exercise performance. Six obese sedentary men with T2D and 7 control participants without diabetes matched for age, sex and body mass index were recruited. Anthropometric characteristics, blood samples, resting cardiac and pulmonary functions and maximal oxygen uptake (VO_2_max) and ventilatory threshold were measured on a first visit. On the four subsequent visits, participants performed step transitions (6 min) of moderate-intensity exercise on an upright cycle ergometer from unloaded pedaling to 80 % of ventilatory threshold. VO_2_ (τVO_2_) and HR (τHR) kinetics were characterized with a mono-exponential model. VO_2_max (27.8±4.0 vs. 27.5±5.3 ml kg^-1^ min^-1^; p=0.95), τVO_2_ (43±6 vs. 43±10 s; p=0.73) and τHR (42±17 vs. 43±13 s; p=0.94) were similar between diabetics and controls respectively. The remaining variables were also similar between groups. These results suggest that well-controlled T2D is not associated with a reduction in VO_2_max or slower τVO_2_ and τHR.

## Introduction

The presence of type 2 diabetes, with or without cardiovascular complications, is often associated with reduced maximal oxygen uptake (VO_2_max) (1-14). Type 2 diabetes also seems to affect submaximal VO_2_ response, however, results are equivocal. A study conducted in older men (65 ± 5 years) with well-controlled type 2 diabetes and long disease duration (> 5 years) (9) reported no difference in VO_2_ kinetics compared to control participants without diabetes, while others reported a slower VO_2_ kinetics in pre- and postmenopausal women (4, 7, 10), adolescents with type 2 diabetes (6) and middle-aged men with type 2 diabetes (10, 15).

The adjustment of heart rate (HR) at the onset of exercise (e.g. HR kinetics), provides additional insights regarding the influence of the central component [central blood flow adjustment and O_2_ delivery (16)] on exercise capacity and, to our knowledge, only a few studies have evaluated HR kinetics in men with type 2 diabetes (9, 10). Recently, O’Connor et al. (15) have demonstrated slowed HR kinetics in older men with type 2 diabetes.

Taken together, these findings support the fact that this metabolic disorder eventually affects negatively numerous human body functions depending on patients’ glycemic control, disease duration as well as the presence of comorbidities due to diabetes. Whether abnormal exercise capacity is the result of type 2 diabetes *per se* or a consequence of its associated comorbidities is not clear, as most of the studies on men have been conducted in patients with type 2 diabetes without optimal glycemic control (9, 10). Evidence of subclinical abnormalities in autonomic (17) and cardiac function (18-21) have been reported in patients with diabetes. Reduced heart rate variability (22), left ventricular diastolic dysfunction (23) and lower lung capacity (24) have all been associated with reduced maximal exercise capacity in patients with type 2 diabetes. Still, the influence of type 2 diabetes *per se* on VO_2_ and HR kinetics in uncomplicated well-controlled middle-aged men remains unknown.

The aim of this study was to evaluate the impact of well-controlled, uncomplicated type 2 diabetes on VO_2_max as well as VO_2_ and HR kinetics in obese sedentary middle-aged type 2 diabetes men compared to control participants matched for sex, age and body mass index. A secondary objective of this study was to provide an integrative view of the impact of type 2 diabetes, including metabolic control, lipid profile, cardiopulmonary function, cardiac structures, and heart rate variability, in relationship to exercise performance. We hypothesized that the presence of well-controlled, uncomplicated type 2 diabetes would be associated with a reduction in VO_2_max but similar VO_2_ and HR kinetics compared to participants without diabetes.

## Methods

### Study population

Six sedentary men with type 2 diabetes and 7 control participants without diabetes matched for age, sex, and body mass index were recruited for this study. This control group, which represents a real life situation considering the impacts of an inactive lifestyle, age and overweight/obesity on human body systems and exercise performance (25-27), strengthen the study design. Type 2 diabetes was diagnosed according to American Diabetes Association criteria (28). All subjects with diabetes were managed with diet, except two who were treated with metformin alone. No participants were using insulin or any cardiovascular drug regimen for the treatment of other diseases/comorbidities. Exclusion criteria for both groups were the documented presence of cardiovascular disease, a documented office blood pressure above 140/90 mmHg and clinically significant end-organ complications related to diabetes, i.e. renal failure (creatinine above normal upper limit), macroalbuminuria, proliferative retinopathy, clinically significant sensitive, motor or autonomic neuropathies as well as the participation to a structured exercise training program. The local ethics committee approved the study, in accordance with the Helsinki declaration, and all participants gave signed informed consent.

### Study design

Participants visited our laboratory on 5 different occasions. Blood profile and a resting echocardiographic variables, pulmonary function, heart rate variability, and VO_2_max were evaluated on the first visit (V_1_). Then, participants performed square-wave transitions from unloaded pedaling to moderate-intensity exercise on four different visits separated by 48 hours (V_2_-V_5_) to determine VO_2_ and HR kinetics.

### Measurements

#### Blood samples

Upon participant arrival at the laboratory (V_1_), a 20-gauge polyethylene catheter was inserted into a forearm vein for blood sampling. Blood samples were drawn at rest after an overnight fasting. Fasting blood glucose was measured using the hexokinase method (Roche Diagnosis, Indianapolis, IN). Glycated hemoglobin was assayed using the ion-exchange high-performance liquid chromatography method (Bio-Rad, Hercules, CA). Serum cholesterol, triglycerides and high-density lipoprotein-cholesterol were analyzed as previously described (21, 29). Low-density lipoprotein-cholesterol was calculated using Friedewald’s formula (30). The cholesterol/high-density lipoprotein ratio was also calculated.

#### Echocardiography

Standard parasternal, short- and long-axis and apical views were performed in accordance with the recommendations of the American Society of Echocardiography (31) with the same observer obtaining all recordings and measurements (Sonos 5500; Hewlet Packard, Andover, Massachusetts) (29). Left atrial volume (LAV) and left ventricular (LV) systolic and diastolic volumes were calculated using the modified Simpson’s method. Left ventricular mass (LVM) and wall thickness were evaluated by M-mode Doppler. LVM was calculated using the following formula (32): LVM (g) = 0.8 × 1.04 [(LVEDD + IVST + PWT)^3^ − (LVEDD)^3^] + 0.6, where LVEDD is the LV end diastolic dimension, IVST the interventricular septal thickness, and PWT the posterior wall thickness. LAV and LVM were indexed for body surface area (33). Ejection fraction was evaluated using the Simpson’s method.

Resting left ventricular diastolic dysfunction (LVDD) was evaluated using standardized criteria (29, 34). First, transmitral pulsed Doppler recordings were obtained to measure the following parameters: peak E velocity in cm/s (peak early transmitral filling velocity during early diastole), peak A velocity in cm/s (peak transmitral atrial filling velocity during late diastole), deceleration time in ms (time elapsed between peak E velocity and the point where the extrapolation of the deceleration slope of the E velocity crosses the zero baseline) and E/A ratio (peak E wave velocity divided by peak A wave velocity). In order to reduce the high filling pressures encountered in the pseudonormalized pattern of LV filling, the same measurements were repeated during phase II of the Valsalva maneuver (29). Tissue Doppler velocities were measured at the mitral annular level of the LV septum, to provide additional information about filling pressure. Early (Ea) and late (Aa) velocities were measured and E/Ea ratio was calculated (35). LVDD was characterized as normal, abnormal relaxation, pseudonormal pattern and restrictive pattern. Subjects had a normal LV diastolic function if a deceleration time between 140-240 msec, an E/A ratio between 1 and 2 and an E/Ea ratio < 8 were present. LVDD was characterized as abnormal relaxation if subjects had a deceleration time > 240 msec and an E/A ratio < 1. LVDD was characterized as a pseudonormal filling pattern if subjects had a deceleration time between 150-240 msec, an E/A ratio between 1 and 2 and an E/Ea ratio > 15 (36), or a reduction in the E/A ratio > 0.5 following the Valsalva maneuver. Finally, LVDD was characterized as a restrictive filling pattern if subjects had a deceleration time < 140 msec and E/A ratio > 2.

#### Pulmonary function tests

Standard pulmonary function tests, including body plethysmography, spirometry and single-breath diffusing capacity of the lung for carbon monoxide [D_LCO_] were performed in participants (37).

#### Heart rate variability

Heart rate variability was derived from a 24-hour Holter monitoring system (Marquette Electronics, Milwaukee, WI) in all participants during normal daily life activity. Heart rate variability derived from 24-hour ambulatory monitoring is reproducible and free of placebo effect (38). Within the 24-hour evaluation, three periods were arbitrarily assessed: 1) 24 hours, 2) daytime period defined as 8:00 AM to 8:00 PM and, 3) night-time period defined as 12:00 AM to 6:00 AM (23). In the frequency domain, power in the low-frequency (0.04 to 0.15 Hz), and high-frequency (0.15 to 0.4 Hz) ranges were calculated. The low frequency/high frequency ratio, considered to be a marker of the ratio of sympathetic to parasympathetic balance, was also determined (39). Using time domain analysis, the standard deviation (SD) of the RR intervals (SDNN), the square root of the mean squared differences of successive RR intervals (rMSSD), and the SD of the average RR intervals were calculated over 5-minute periods (SDANN) and the average of the SD of RR intervals for all 5-minute periods (ASDNN) were determined. pNN50 is the proportion of interval differences of successive NN intervals >50 ms. rMSSD and pNN50 are indices of parasympathetic modulation. NN intervals are the normal-to-normal intervals that include all intervals between adjacent QRS complexes resulting from sinus node depolarizations in the entire 24-hour electrocardiogram recording. The complete signal was carefully edited using visual checks and manual corrections of individual RR intervals and QRS complex classifications.

#### Blood pressure

Following 15 minutes of quiet rest in a supine position, resting arterial blood pressure was measured with the participants seated using an automated sphygmomanometer with headphone circuit option before the maximal exercise protocol (Model 412, Quinton Instrument Co., Bothell, WA, USA).

#### Maximal exercise protocol

VO_2_max was evaluated using a ramp incremental exercise protocol of 20 watts/min following a warm-up period of 1 minute of unloaded pedaling, performed on an electromagnetically braked upright cycle ergometer (Lode Corival, Lode, Groningen, Netherlands) at a pedaling rate of 50 to 70 rpm. Expired air was continuously collected for the determination of pulmonary VO_2_, carbon dioxide production (VCO_2_), pulmonary ventilation (V_E_) and the respiratory exchange ratio (RER) (VCO_2_/VO_2_), on a breath-by-breath basis (Medgraphics, CPX Ultima, St Paul, MN). Heart rate was measured using electrocardiographic monitoring during the test. Participants exercised until volitional exhaustion. VO_2_max was defined as the mean VO_2_ recorded in the last 15 seconds of the ramp protocol concurrent with a RER ≥ 1.15. The ventilatory threshold was evaluated with the V-slope method (40). The exercise protocol was always performed at the same time of the day at a room temperature of 19 °C.

#### Square-wave exercise protocol

Each participant performed four square-wave exercise protocols, from unloaded pedaling to 80% ventilatory threshold, on four different visits separated by at least 48 hours. Pulmonary gas exchange and HR were continuously monitored during exercise.

### Data analysis and statistics

#### VO_2_ and HR kinetics analysis

Breath-by-breath data were filtered and linearly interpolated to provide second-by-second values, then time-aligned to the onset of exercise and ensemble-averaged into 5-s bins. The phase 1 response (approximately 20 seconds) was omitted (41) and a mono-exponential equation was used to fit the data (Origin software, OriginLab, Northampton, MA, USA):

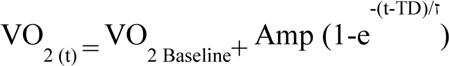

where VO_2(t)_ represents VO_2_ as a function of time *t*; VO_2baseline_ represents the mean VO_2_ in the baseline period (unloaded pedaling); *Amp* is the amplitude or the difference between the baseline and steady-state VO_2_, *TD* is the time delay before the onset of exercise and τVO_2_, the phase 2 time constant, representing the time required to reach 63 % of the steady-state response. The same equation was used for the HR kinetics analysis, starting at time 0. Accordingly, there was not time delay incorporated into the model employed to describe HR kinetics.

#### Statistical analysis

The Mann-Whitney Rank Sum test was used to compare variables between groups. All data is presented as mean ± standard deviation unless otherwise specified. A p value < 0.05 was considered statistically significant. A sample size calculation was based on previous work that studied VO_2_ kinetics (7, 42) in women with type 2 diabetes. Accordingly, 8 participants per group were necessary in order to report a statistically significant difference of 15±10 sec in τVO_2_ between our two groups with a power of 80% and a p<0.05.

## Results

Anthropometric characteristics, resting hemodynamics, metabolic profile (Table 1), cardiac structures, baseline LV systolic function, heart rate variability (Table 2) and pulmonary function (Table 3) were all similar between groups. A higher peak early E wave velocity during the Valsalva maneuver (56±4 vs. 47±5 ms; p<0.05) and lower tissue Doppler septal early velocities (7.6±1.3 vs. 10.9±2.9 cm s^-1^; p<0.05) were observed in diabetics vs. controls. Three participants with diabetes and four control participants had LVDD. There was a trend toward higher LV filling pressure (measured by the septal E/Ea ratio) (9.1±2.8 vs. 7.0±1.6; p=0.06) in diabetics vs. controls.

**Table 1.** Baseline characteristics, blood profile and resting hemodynamics in participants with type 2 diabetes compared to control participants. Values are means **±** SD; *Data presented as median (minimum-maximum) BMI: Body mass index; Hb: hemoglobin HR: Heart rate; SBP: Systolic blood pressure; DPB: Diastolic blood pressure; FBG: Fasting blood glucose; HbA_1c_: Glycated hemoglobin; HDL-C: High-density lipoprotein cholesterol; LDL-C: Low-density lipoprotein cholesterol

|  | Diabetes group | Control group | p |
| --- | --- | --- | --- |
| N | 6 | 7 | - |
| Age (yrs) | $58 \pm 2$ | $54 \pm 9$ | 0.43 |
| Height (m) | $1.76 \pm 0.07$ | $1.77 \pm 0.03$ | 1.00 |
| Body weight (kg) | $90 \pm 17$ | $97 \pm 15$ | 0.29 |
| BMI (kg/m<sup>2</sup>) | $29 \pm 4$ | $31 \pm 5$ | 0.53 |
| Diabetes duration (month)* | 50 (4-138) | - | - |
| FBG (mmol l<sup>-1</sup>) | $6.3 \pm 2$ | $5.1 \pm 0.4$ | 0.19 |
| HbA<sub>1c</sub> (%) | $6.2 \pm 0.8$ | $5.7 \pm 0.3$ | 0.25 |
| HR (bpm) | $68 \pm 9$ | $66 \pm 9$ | 0.52 |
| SBP (mmHg) | $133 \pm 9$ | $143 \pm 10$ | 0.12 |
| DPB (mmHg) | $87 \pm 7$ | $88 \pm 7$ | 0.77 |
| Cholesterol (mmol l<sup>-1</sup>) | $5.3 \pm 1.2$ | $5.3 \pm 0.6$ | 0.72 |
| Triglycerides (mmol l<sup>-1</sup>) | $1.7 \pm 0.9$ | $1.8 \pm 0.8$ | 1.0 |
| HDL-C (mmol l<sup>-1</sup>) | $1.3 \pm 0.2$ | $1.2 \pm 0.3$ | 0.14 |
| LDL-C (mmol l<sup>-1</sup>) | $3.1 \pm 1.0$ | $3.4 \pm 0.6$ | 0.73 |
| Total-Cholesterol/HDL | $4.1 \pm 1.2$ | $4.9 \pm 1.2$ | 0.29 |

**Table 2.**
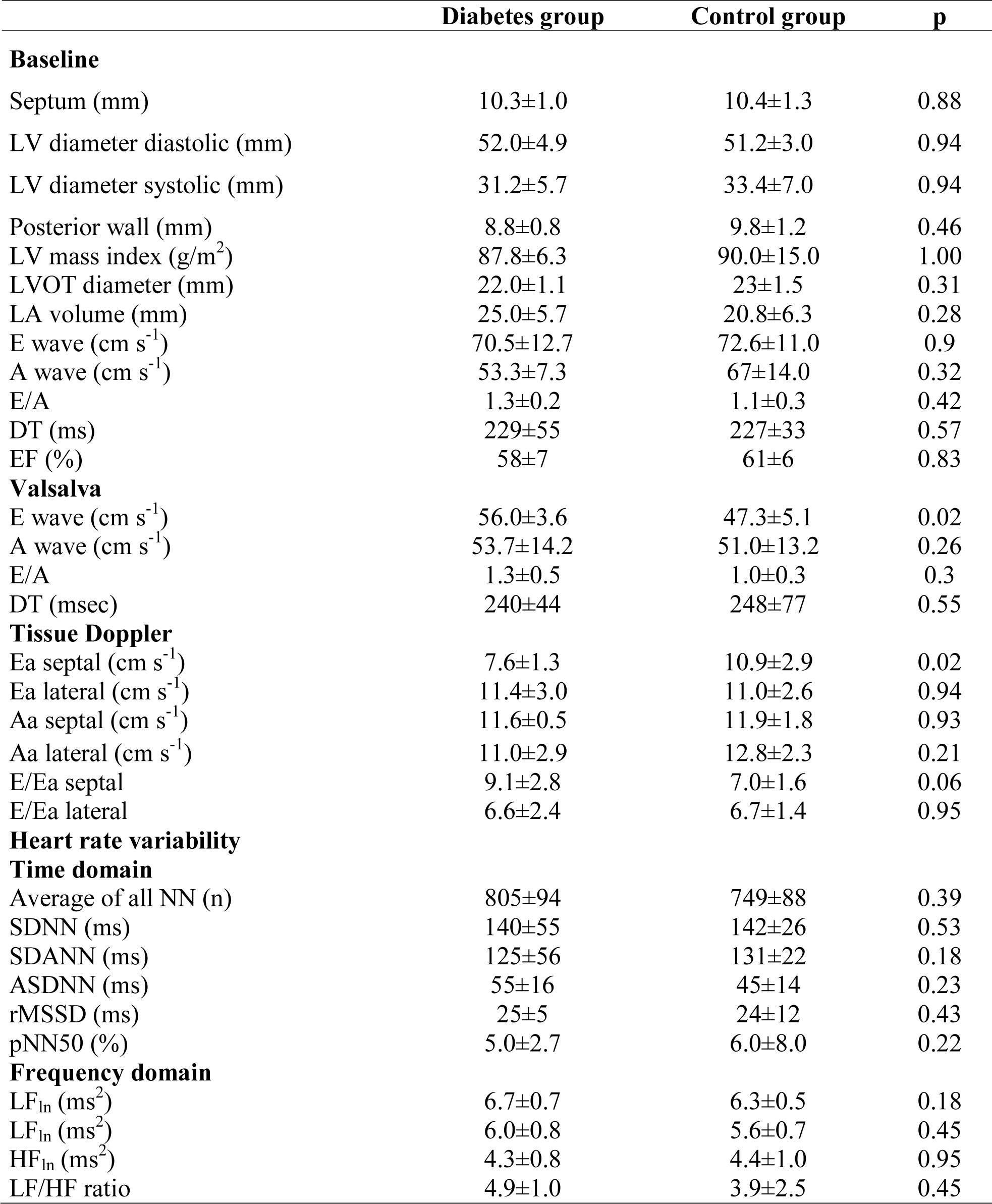
Cardiac structure and function and heart rate variability in participants with type 2 diabetes compared to control participants. Values are means **±** SD; LV: Left ventricular; LA; Left atrial; LVOT: Left ventricular outflow track; E: Mitral early diastolic velocity; A: Mitral late diastolic velocity; DT: Deceleration time; Ea: Mitral annulus early diastolic velocity; Aa: Mitral annulus late diastolic velocity; NN: Normal to normal interval; SDNN: Standard deviation of all NN intervals; SDANN: SD of the average NN intervals for all 5-min segments; ASDNN: average of the standard deviationof NN intervals for all 5-min segments; rMSSD: Square root of the mean of the squared differences between adjacent NN intervals; pNN50: NN50 count divided by the total number of all NN intervals; VLF: Very low frequency; LF: Low frequency; HF: High frequency; LF/HF ratio: Low frequency to high frequency ratio

|  | Diabetes group | Control group | p |
| --- | --- | --- | --- |
| **Baseline** |  |  |  |
| Septum (mm) | 10.3±1.0 | 10.4±1.3 | 0.88 |
| LV diameter diastolic (mm) | 52.0±4.9 | 51.2±3.0 | 0.94 |
| LV diameter systolic (mm) | 31.2±5.7 | 33.4±7.0 | 0.94 |
| Posterior wall (mm) | 8.8±0.8 | 9.8±1.2 | 0.46 |
| LV mass index (g/m<sup>2</sup>) | 87.8±6.3 | 90.0±15.0 | 1.00 |
| LVOT diameter (mm) | 22.0±1.1 | 23±1.5 | 0.31 |
| LA volume (mm) | 25.0±5.7 | 20.8±6.3 | 0.28 |
| E wave (cm s<sup>-1</sup>) | 70.5±12.7 | 72.6±11.0 | 0.9 |
| A wave (cm s<sup>-1</sup>) | 53.3±7.3 | 67±14.0 | 0.32 |
| E/A | 1.3±0.2 | 1.1±0.3 | 0.42 |
| DT (ms) | 229±55 | 227±33 | 0.57 |
| EF (%) | 58±7 | 61±6 | 0.83 |
| **Valsalva** |  |  |  |
| E wave (cm s<sup>-1</sup>) | 56.0±3.6 | 47.3±5.1 | 0.02 |
| A wave (cm s<sup>-1</sup>) | 53.7±14.2 | 51.0±13.2 | 0.26 |
| E/A | 1.3±0.5 | 1.0±0.3 | 0.3 |
| DT (msec) | 240±44 | 248±77 | 0.55 |
| **Tissue Doppler** |  |  |  |
| Ea septal (cm s<sup>-1</sup>) | 7.6±1.3 | 10.9±2.9 | 0.02 |
| Ea lateral (cm s<sup>-1</sup>) | 11.4±3.0 | 11.0±2.6 | 0.94 |
| Aa septal (cm s<sup>-1</sup>) | 11.6±0.5 | 11.9±1.8 | 0.93 |
| Aa lateral (cm s<sup>-1</sup>) | 11.0±2.9 | 12.8±2.3 | 0.21 |
| E/Ea septal | 9.1±2.8 | 7.0±1.6 | 0.06 |
| E/Ea lateral | 6.6±2.4 | 6.7±1.4 | 0.95 |
| **Heart rate variability** |  |  |  |
| **Time domain** |  |  |  |
| Average of all NN (n) | 805±94 | 749±88 | 0.39 |
| SDNN (ms) | 140±55 | 142±26 | 0.53 |
| SDANN (ms) | 125±56 | 131±22 | 0.18 |
| ASDNN (ms) | 55±16 | 45±14 | 0.23 |
| rMSSD (ms) | 25±5 | 24±12 | 0.43 |
| pNN50 (%) | 5.0±2.7 | 6.0±8.0 | 0.22 |
| **Frequency domain** |  |  |  |
| LF<sub>ln</sub> (ms<sup>2</sup>) | 6.7±0.7 | 6.3±0.5 | 0.18 |
| LF<sub>ln</sub> (ms<sup>2</sup>) | 6.0±0.8 | 5.6±0.7 | 0.45 |
| HF<sub>ln</sub> (ms<sup>2</sup>) | 4.3±0.8 | 4.4±1.0 | 0.95 |
| LF/HF ratio | 4.9±1.0 | 3.9±2.5 | 0.45 |

**Table 3.** Pulmonary function at rest and maximal exercise parameters in participants with diabetes and control participants. Values are means **±** SD; FEV1: Forced expiratory volume-second; DLCO: Diffusion capacity to carbon dioxide; VO_2_: Oxygen uptake; HR: Heart rate SBP: Systolic blood pressure; DBP: Diastolic blood pressure; V_E_: Minute-ventilation; BF: Breathing frequency; RER: Respiratory exchange ratio

|  | Diabetes group | Control group | p |
| --- | --- | --- | --- |
| **Pulmonary function** |  |  |  |
| Total lung capacity (l) | 6.9±0.8 | 7.4±0.8 | 0.8 |
| Vital capacity (l) | 5.2±0.5 | 5.5±0.7 | 0.4 |
| Forced vital capacity (l) | 5.1±0.6 | 5.3±0.9 | 0.8 |
| FEV<sub>1</sub> (l) | 3.7±0.4 | 3.95±0.70 | 0.5 |
| Residual volume (l) | 1.7±0.4 | 1.9±0.6 | 0.9 |
| Inspiratory capacity (l) | 3.8±0.9 | 3.95±0.4 | 0.7 |
| DLCO (ml mm Hg<sup>-1</sup> min<sup>-1</sup>) | 30.6±6.3 | 27.6±2.3 | 0.3 |
| **Exercise capacity** |  |  |  |
| VO<sub>2</sub>max (l min<sup>-1</sup>) | 2.59±0.44 | 2.62±0.33 | 0.95 |
| VO<sub>2</sub>max (ml·kg<sup>-1</sup>·min<sup>-1</sup>) | 27.8±4.0 | 27.5±5.3 | 1.00 |
| Work rate at VO<sub>2</sub> max (W) | 217±24 | 217±22 | 1.00 |
| Ventilatory threshold (L/min) | 1.6±0.2 | 1.3±0.2 | 0.10 |
| Maximal HR (beat min<sup>-1</sup>) | 164±17 | 165±18 | 1.00 |
| Maximal SBP (mmHg) | 210±16 | 231±18 | 0.07 |
| Maximal DBP (mmHg) | 84±11 | 86±9 | 0.5 |
| Maximal V<sub>E</sub> (l min<sup>-1</sup>) | 120±35 | 106±23 | 0.7 |
| RER | 1.2±0.1 | 1.3±0.1 | 0.4 |

VO_2_max and cardiopulmonary responses at maximal exercise were similar between groups (Table 3). Both τVO_2_ (43±6 vs. 43±10 s; p=0.73; Figure 1) and τHR (42±17 vs. 43±13 s; p=0.94; Figure 2) were similar between diabetics and control participants. Heart rate amplitude was greater in diabetics (p<0.05; Table 4). However, the other VO_2_ and HR kinetics variables were similar between groups (Table 4).

**Table 4.**
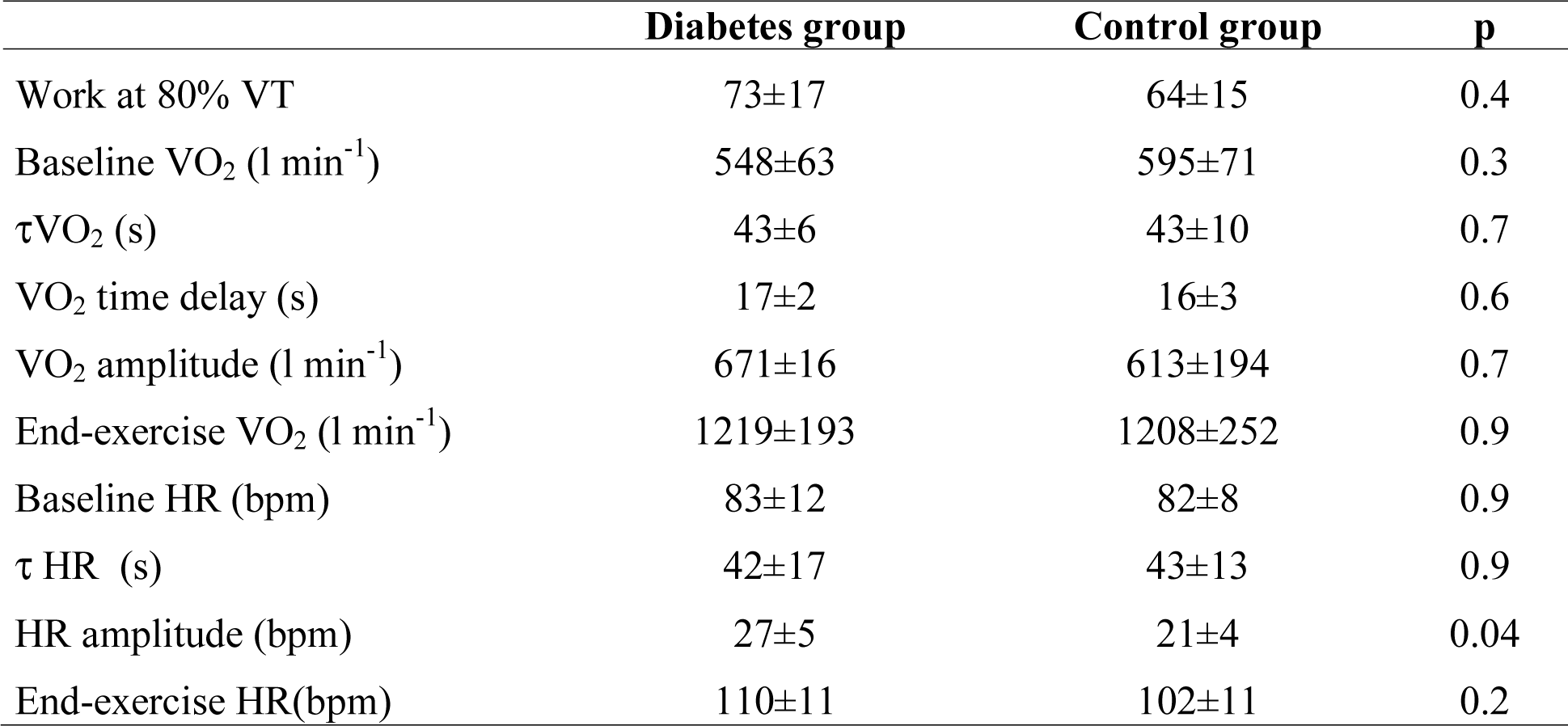
VO_2_ and heart rate kinetics during exercise in participants with type 2 diabetes compared to control participants. Values are means **±** SD; HR: Heart rate; VO_2_: Oxygen uptake; VT: Ventilatory threshold

**Figure 1.**
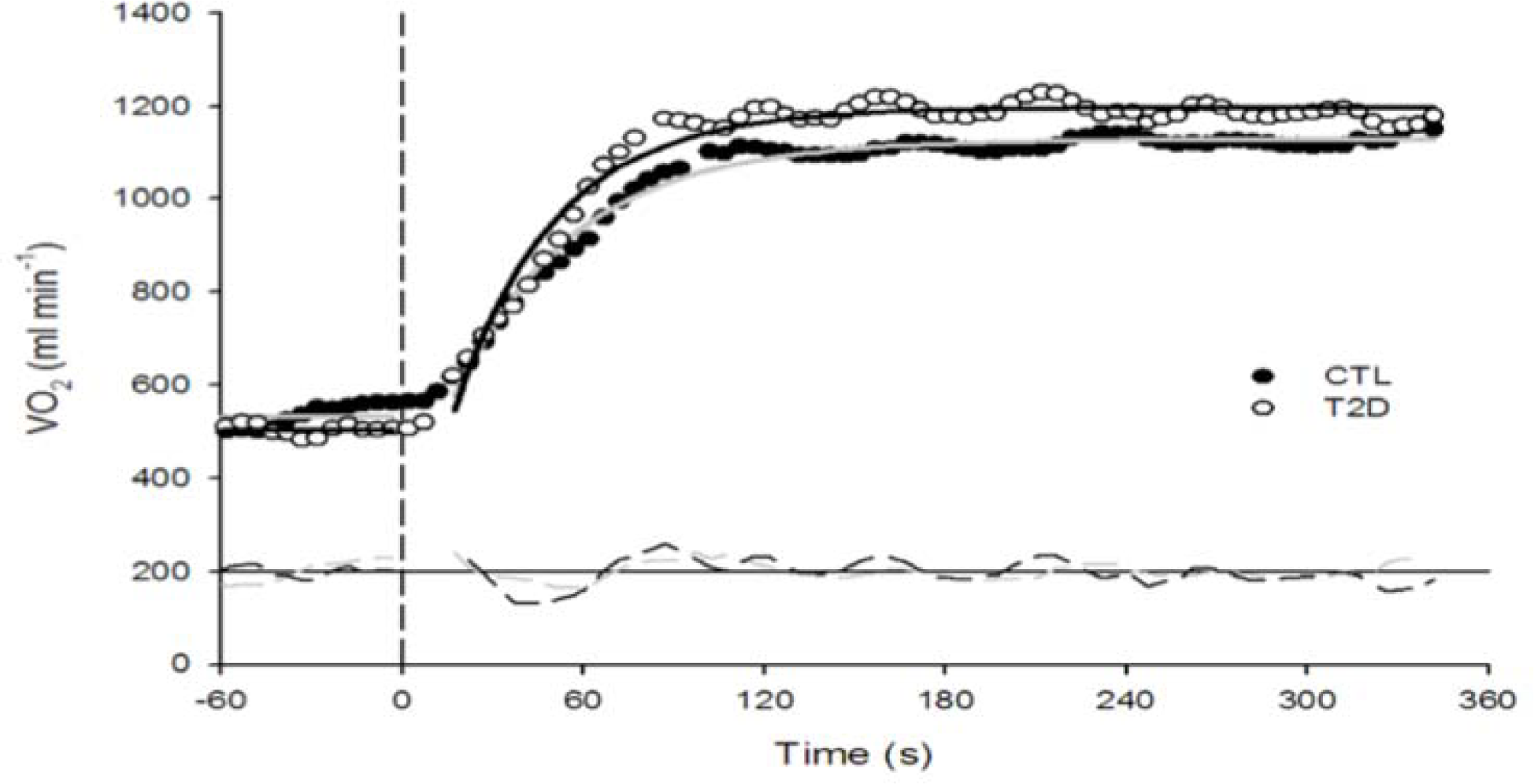
VO_2_ kinetic response of a representative participant with type 2 diabetes vs. a control participant. Full circles represent single breath by control participant and overall response is characterized by the grey line. Empty circles represent single breath by participants with type 2 diabetes while the overall response is characterized by the black line. No significant difference between groups in the phase 2 VO_2_ kinetic response.

**Figure 2.**
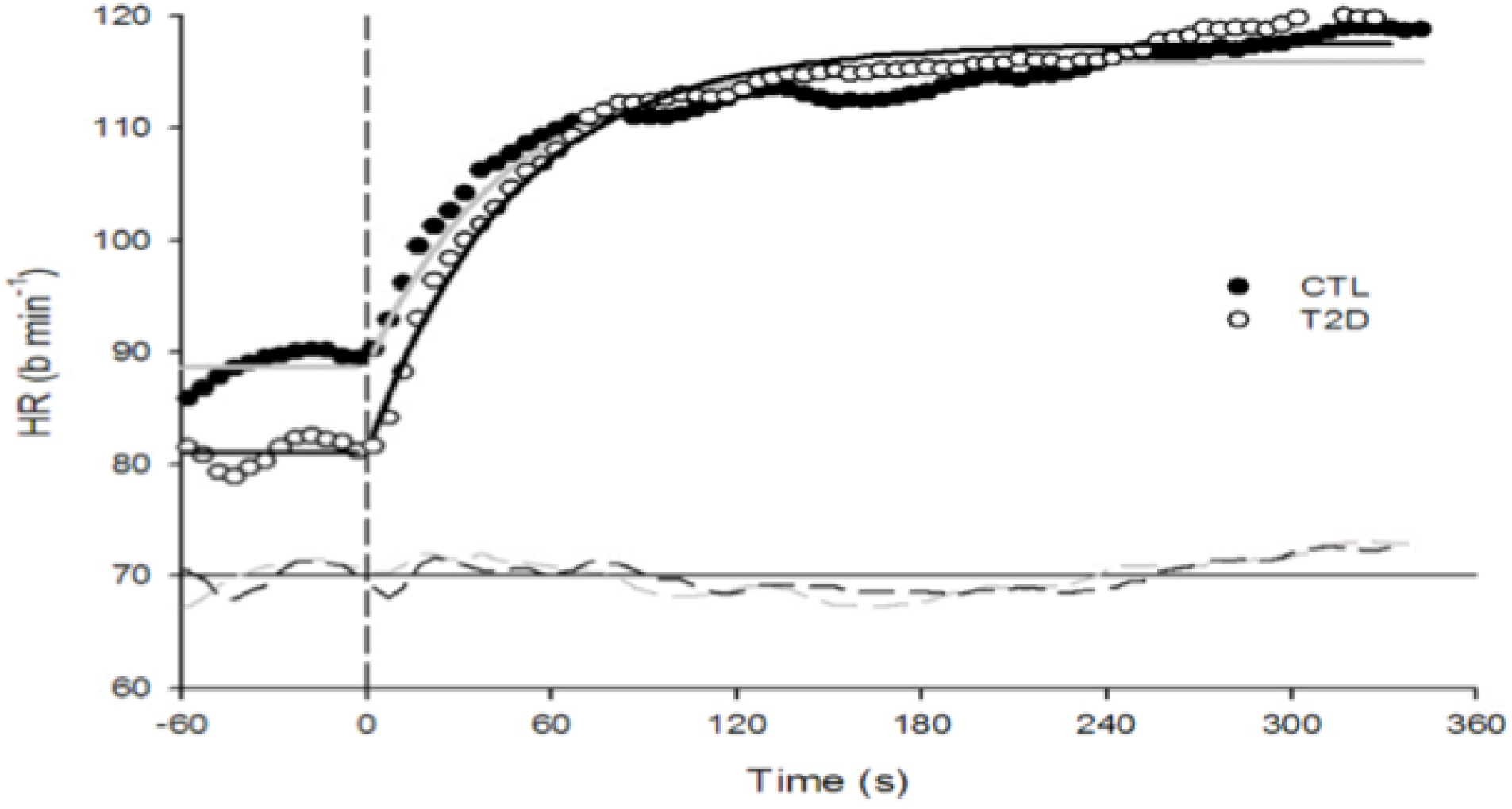
HR kinetic response of a representative participant with type 2 diabetes vs. a control participant. Full circles represent single breath by control participants and overall response is characterized by the grey line. Empty circles represent single breath by participants with type 2 diabetes while the overall response is characterized by the black line. No significant difference between groups in the phase 2 HR kinetic response.

## Discussion

The results of this study suggest that obese sedentary men with well-controlled type 2 diabetes without clinical complications or comorbidities do not show a reduction in VO_2_max or slower VO_2_ and HR kinetics compared to participants without diabetes matched for sex, age and body mass index. In addition, diabetics and control participants had similar metabolic profile and resting cardiorespiratory function, which may explain the similar submaximal and maximal exercise capacity responses reported in this study.

Type 2 diabetes has been associated with reduced maximal exercise performance (2, 4-11, 14). However, the exact underlying pathophysiological mechanisms are still ambiguous. In our study including well-matched participants, no difference in VO_2_max was observed between groups, in contrast to previous reports. It is noteworthy that in most of the studies that reported a reduction in VO_2_max in patients with diabetes, other important variables may have contributed to the reduced VO_2_max, such as poor glycemic control (14, 43, 44), presence of LVDD (20, 21), reduced heart rate variability (20, 45), impaired lung diffusion capacity or early signs of cardiac impairment altering pumping capacity (46, 47) and the presence of comorbidities associated with type 2 diabetes (13). The presence of type 2 diabetes did not add additional burden to the majority of variables related to resting cardiopulmonary function in the present study. Since the cardiopulmonary function has an important influence on exercise performance in healthy individuals (47), the absence of a deleterious impact of well-controlled, uncomplicated type 2 diabetes on VO_2_max in this study is not surprising. It is noteworthy that our participants with type 2 diabetes were treated with diet (n=4) or metformin alone (n=2) reinforcing the fact that their diabetes was not longstanding which may have required a more aggressive pharmacological intervention.

In addition, type 2 diabetes is usually associated with altered LV diastolic function (18, 45, 48, 49). The negative impact of LVDD on submaximal exercise responses (8) and maximal exercise capacity (21) has been reported. Although a trend toward higher LV filling pressure was observed in diabetics vs. controls, a similar number of participants in both groups had LVDD, which could partly contribute to the comparable maximal exercise capacity between the two groups. What is still ambiguous pertaining to LVDD in diabetics is whether its appearance coincides with the development of diabetes or occurs earlier, since several variables such as blood pressure, age, sedentary lifestyle, obesity, hyperinsulinemia and LV systolic failure may modulate the development of LVDD (26, 27, 50). Accordingly, the results of this study do not exclude that LVDD has a negative influence on exercise capacity in patients with type 2 diabetes (51). Further research is necessary to isolate the influence of LVDD on VO_2_max in these patients.

The evaluation of VO_2_ and HR kinetics provides supplemental mechanistic information on how a clinical population, in this case well-characterized men with well-controlled type 2 diabetes with short known disease duration, adapt to a transition from low to moderate intensity exercise. Compared to VO_2_max, VO_2_ and HR kinetics provide a more accurate evaluation of how participants adapt to everyday life activity, considering that they rarely perform activities at maximal exercise intensity. As VO_2_ kinetics represents the efficiency of the cardiovascular and metabolic systems to respond to changes in demand, a slower VO_2_ or HR adjustment at the onset of exercise is associated with a lower functional capacity and harder perceived effort when performing regular activities (16). A slower response may be the result of impaired O_2_ delivery, inappropriate distribution of O_2_ to the working muscles, other muscular factors (metabolic inertia), or more likely, a combination of these factors (16).

In our study, no difference in VO_2_ kinetics was observed between groups. This contrasts with previous studies conducted in patients with type 2 diabetes, in which slower VO_2_ kinetics has been reported (4, 6, 7, 15). Factors that could have contributed to the slower VO_2_ kinetics reported in previous studies include non-optimal short and long-term glycemic control, comparison with a lean control group, reduced heart rate variability and/or other signs of cardiac impairment, such as the presence of LVDD in diabetics. Of note, LVDD was not associated with slower VO_2_ kinetics in this study, but this could be related to our small sample size. However, our results support those of Wilkerson et al. (9) who reported similar VO_2_ kinetics between older men with diabetes of longer disease duration vs. control subjects. This similar VO_2_ adjustment was attributed to altered blood flow compensated by adaptive mechanisms with long-disease duration, such as O_2_ extraction (52). Interestingly, the slowed VO_2_ kinetics reported in uncomplicated type 2 diabetics could attain a plateau early following the onset of the disease, without a further detrimental impact of aging (15).

The evaluation of HR kinetics provides a measure of central blood flow adjustment, and O_2_ delivery, at the onset of moderate exercise (16). Previous studies having investigated HR kinetics in type 2 diabetics provided equivocal observations. The adjustment of HR seems slower in pre and post-menopausal women (7, 10), as well as in older men with type 2 diabetes with longer disease duration (9) or with sub-optimal glycemic control (10). The difference between those results and ours may be attributable to the shorter disease duration of our cohort (< 5 years), the mean age of our participants and the optimal short and long-term glycemic control. In addition, diabetics involved in this study were treated with diet or metformin alone, compared to more advanced treatment (10), which often reflects a more advanced form of diabetes or the presence of complications or comorbidities. To our knowledge, the present study is the first to show that well-characterized optimally-controlled men with type 2 diabetes and of a relatively short known disease duration (< 5 years) do not have slower HR kinetics compared to control subjects. In addition, it appears that the slowed HR kinetics may be influenced to a greater extent by other variables, such as age, fitness or the presence of comorbidities rather than the disease processes, at least in the early stage of diabetes.

This study has limitations that need to be discussed. We acknowledge that the number of participants in the present study was relatively low, only composed of men and the study did not include a lean control group. These factors limit the generalization of our results. Although 8 participants per group would have been necessary to report a statistically significant difference of 15±10 sec in τVO_2_ between our groups based on previous reported data (7, 42), we did not reach such a sample size (diabetic group: n=6 and control group: n=7). However, it is highly unlikely that τVO_2_ would have been different with the addition of 2 participants in the diabetic group and 1 participant in the control group taking into consideration that the difference in τVO_2_ between groups was far from the ∼15 sec reported in the literature. By study design, participants in both groups had similar age and body mass index. They also had similar glycemic control, lipid profile, resting blood pressure, pulmonary function as well as cardiac autonomic system modulation. The majority of participants with diabetes were recently diagnosed, and all of them were well-controlled, while control participants may not be considered as completely “healthy” participants, considering their age, the presence of obesity in a majority of participants and the fact that they were physically inactive. We consider that the similarities observed between our groups is a strength of this study which permit to assess the impact of type 2 diabetes *per se*, and not the related complications, on submaximal and maximal exercise performance.

The number of square-wave exercise protocols used to model VO_2_ and HR kinetics is also a strength of this study. Indeed, at least 3 transitions seem necessary to get an adequate signal-to-noise ratio (53) and previous studies that investigated VO_2_ or HR kinetics in patients with type 2 diabetes did not all reach that prerequisite, which considerably reduces the precision in the modeling of the underlying physiologic responses during an exercise transition. Another important issue is that we allowed 48 hours between each exercise trial for the evaluation of VO_2_ and HR kinetics, to ensure that insulin sensitivity-induced exercise returned to baseline value (54), and that fatigue would not interfere with the results, considering the low level of physical fitness of our population. Finally, these results warrant further studies investigating the influence of well-controlled, uncomplicated type 2 diabetes on exercise performance. Still, these results have potential clinical implications. If the presence of type 2 diabetes eventually leads to a lower capacity to perform exercise (14), the present study suggests that well-controlled type 2 diabetes is not always associated with a reduction in submaximal (VO_2_ and HR kinetics) or maximal exercise performance (VO_2_ max). A more likely interpretation of these results may be that exercise performance is already impaired prior to the diagnosis of type 2 diabetes, as observed in participants with metabolic syndrome and healthy first-degree relatives of patients with type 2 diabetes, in whom subclinical metabolic and/or cardiovascular abnormalities are present (55-57). In the eventuality that these findings are supported in further studies, it will reinforce the importance of strong early therapeutic actions in individuals prone to develop type 2 diabetes in order to delay the appearance of this metabolic disorder and preserve the individual’s exercise capacity and quality of life.

In conclusion, the findings from this study suggest that well-controlled type 2 diabetes does not necessarily result in a reduction of VO_2_ max and a slowing of VO_2_/HR kinetics over and above what can be expected in obese and sedentary individuals.

## Acknowledgments

Paul Poirier is a senior clinician-scientist of the *Fonds de recherche du Québec - Santé (FRQS)*. Annie Ferland was supported by the Canadian Institutes of Health Research (CIHR). This study has been supported in part by the Canadian Diabetes Association. Patrice Brassard is a Junior 1 Research Scholar of the *Fonds de recherche du Québec - Santé (FRQS)*.

## Conflict of interest

All authors declare that they have no conflict of interest related to the present study.

